# Glycan size and attachment site location affect electron transfer dissociation (ETD) fragmentation and automated glycopeptide identification

**DOI:** 10.1101/676288

**Authors:** Kathirvel Alagesan, Hannes Hinneburg, Peter H Seeberger, Daniel Varón Silva, Daniel Kolarich

**Author notes:** **Corresponding Author:** A/Prof. Daniel Kolarich, Institute for Glycomics, Building G26, Griffith University, Gold Coast campus, Southport, 4222 QLD, Australia.

## Abstract

We used a small synthetic glycopeptide library to systematically evaluate the effect of glycosylation site location and glycan size on the efficiency of ETD MS/MS fragmentation and subsequent automated identification. Understanding how the physico-chemical properties of glycopeptides influence glycopeptide fragmentation allows for optimizing fragmentation conditions and software assisted data analyses, which rely on informative fragmentation spectra for subsequent data processing to identify glycopeptides. Often, mis-assignment of glycopeptides occurs due to uncertainties such as failure to produce sufficient peptide backbone fragment ions. Our synthetic glycopeptide library contained glycopeptides differing in glycosylation site position within the peptide as well as glycan size (from the pentasaccharide *N*-glycan core to fully sialylated, biantennary *N*-glycans). Different software solutions such as SEQUEST and Amanda were compared for ETD glycopeptide identification. We found that all, glycan size, glycosylation site position within a glycopeptide and individual precursor *m/z* significantly impacted the number and quality of assignable glycopeptide backbone fragments, and thus the likelihood to be correctly identified in software assisted data analyses.

## INTRODUCTION

In mass spectrometry-based glycoprotein characterisation, tandem mass spectrometry (MS/MS) is one of the most powerful tools available. In the analyses of glycopeptides it can provide compositional and structural information about glycans and peptides. Combination of the complementary fragmentation techniques such as electron capture dissociation (ECD) and electron transfer dissociation (ETD) with collision induced dissociation (CID) provides the opportunity to characterise both glycans and their site of attachment within a single MS/MS experiment [1, 2]. For glycopeptides, CID low-energy vibrational activation results in the preferential fragmentation of the carbohydrate moiety, usually with little or no peptide backbone cleavage. High-energy collision dissociation (HCD) allows cleavage of both peptide bonds as well the glycosidic ones, which provides information on peptide sequence and glycan structure within a single experiment. Under optimal collision energy settings, HCD fragmentation of glycopeptides results in distinct Y1 ions (peptide+GlcNAc in the case of *N*-glycans) allowing effective glycopeptide identification [3, 4]. ECD and ETD fragmentation, however, predominantly produce c’ ions and z^.^ radical ions resulting from the cleavage of the N-Cα bond within a peptide [5]. During this type of fragmentation, post-translational modifications (PTMs) still remain intact and attached to the amino acid, thus allowing identification of the site of modification within a peptide sequence [6, 7]. However, some technical and analytical challenges need to be overcome in ETD/ECD glycopeptide fragmentation to make this an even more effective technique, especially if larger modifications such as *N*-glycans are to be reliably analysed by this approach [1, 8]. We used a small library of synthetic *N*-glycopeptides to systematically investigate ETD fragmentation of *N*-glycopeptides to understand the various parameters influencing ETD glycopeptide fragmentation. We also evaluated the potential of automated glycopeptide identification using standard proteomics data analysis software where the respective PTMs are treated as variable modifications.

## MATERIAL AND METHODS

If not otherwise stated, all materials were purchased in the highest possible quality from Sigma-Aldrich (St. Louis, MO, USA).

## GLYCOPEPTIDE SYNTHESIS

All glycopeptides were synthesised by solid phase peptide synthesis (SPPS) using previously reported fluorenylmethoxycarbonyl (Fmoc) protocols [9, 10] (Table 1). Glycopeptides were manually synthesised using a commercially available Wang ChemMatrix^®^ (Sigma-Aldrich) in 5-mL or 10-mL disposable polypropylene syringes with a bottom filter. Sialic acid residues were selectively protected by esterification with benzyl bromide prior their use in SPPS [9, 11-13]. The coupling of the glycosylated Asn building blocks was performed as described by Unverzagt and co-workers [14]. The peptide sequence was inspired by a tryptic glycopeptide present in human Protein C [15] (Uniprot Entry: P04070, ^284^EVFVHPNYSK^293^) and human Immunoglobulin G (IgG) 1-4. Variations of the sequence were produced by altering the *N*-glycosylation sequon towards the N- and C-terminus of the glycopeptide.

**Table 1:**
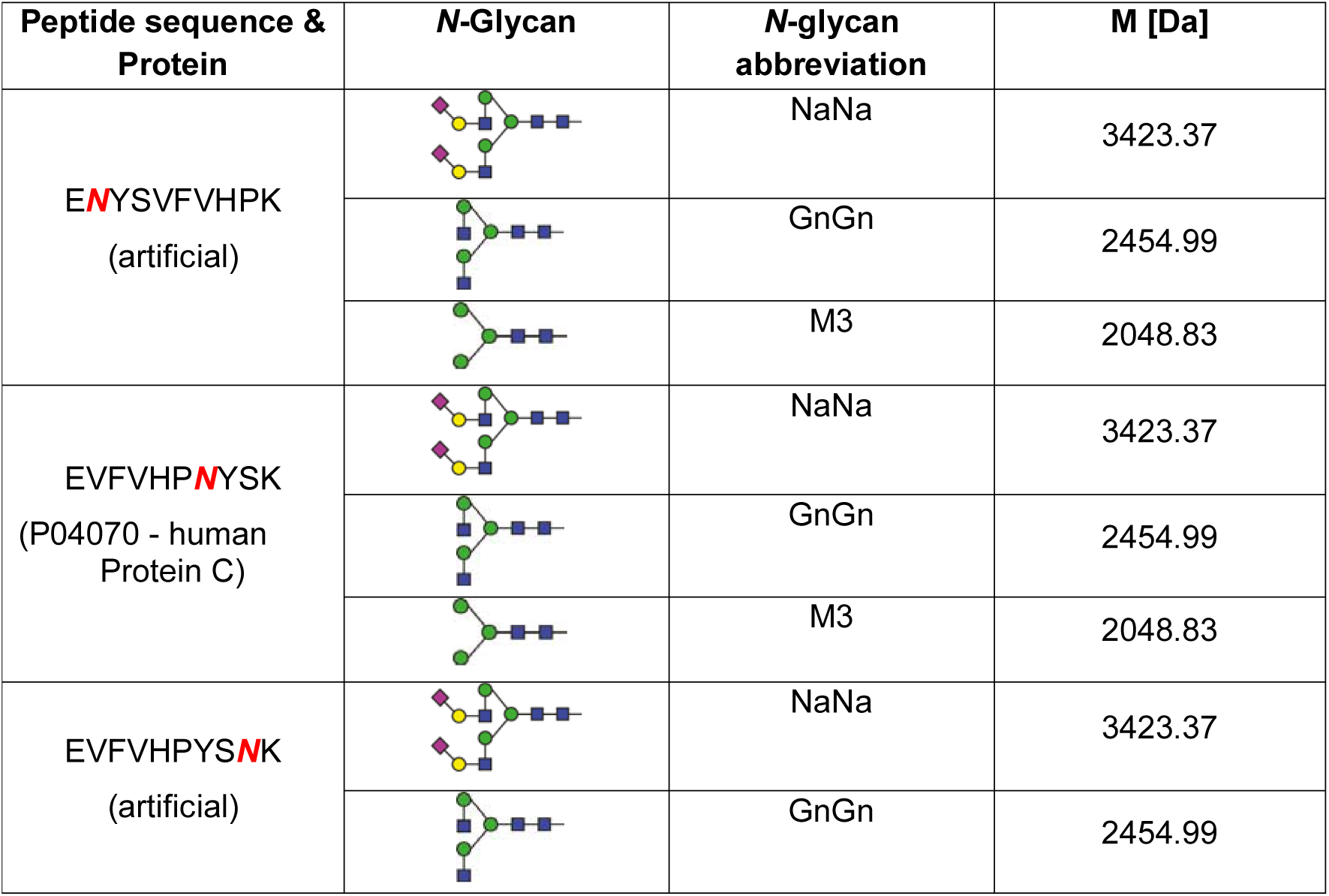
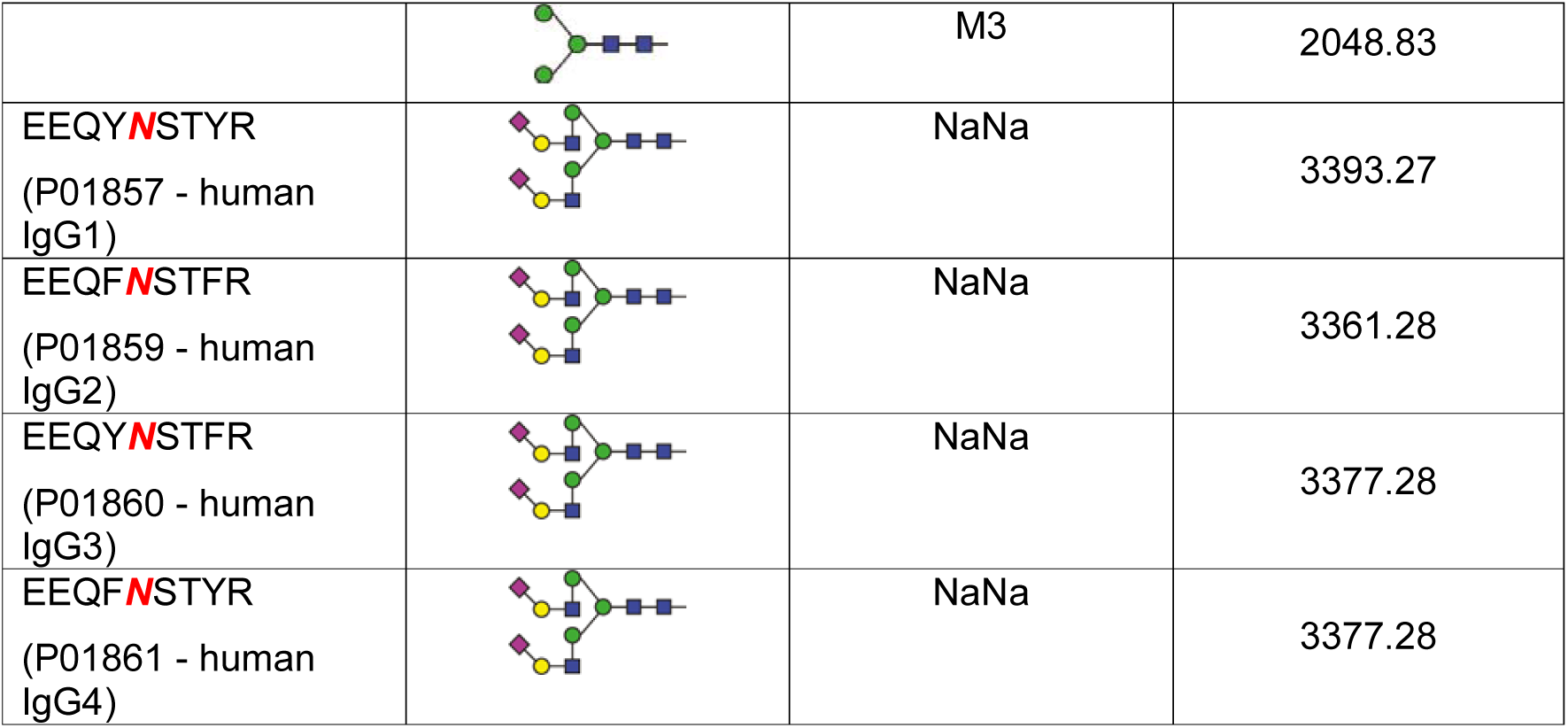
Synthetic N-glycopeptide library used to evaluate the influence of glycan size and precursor m/z on ETD fragmentation

### Mass Spectrometry LC MS/MS

NanoLC-ESI-MS/MS analyses were performed on an Ultimate 3000 RSLC-nano system (Dionex/Thermo Scientific, Sunnyvale, CA) coupled to an amaZon speed ETD ion trap mass spectrometer (IT-MS) operating in positive-ion mode and equipped with the acetonitrile supplying CaptiveSpray nanoBooster™ (both Bruker Daltonics, Bremen, Germany). Briefly, the mass spectrometer was set to perform either CID or ETD on the selected precursor *m/z*. An *m/z* range from 400-1600 Da was used for data dependent precursor scanning. The MS data was recorded using the instrument’s “enhanced resolution mode”. MS/MS data was acquired in “ultra-mode” over an *m/z* range from 100-2000. A detailed overview on the LC-MS parameters is provided in supplementary Table S1 following MIRAGE [16] and MIPAE [17] recommendations. Samples of 3□µl (150□fmol of each glycopeptide) were loaded onto a short C18 column (Acclaim PepMap□100, 100□µm□×□2□cm, 5□µm, 100□Å, Dionex, Part of Thermo Fisher, Germany) and eluted isocratically with 35% buffer B (ACN/0.1% FA) as described previously [12].

Data analysis was performed using DataAnalaysis 4.2 (Bruker Daltonics) and automated glycopeptide identification was performed using Proteome Discoverer 2.1 using SEQUEST and Amanda search engines. Details on the used parameter settings are provided in supplementary Table S2.

## RESULTS AND DISCUSSION

### Glycan size and precursor m/z influence ETD glycopeptide fragmentation

The charge state and *m/z* range of the selected precursor had a considerable impact on the ETD-fragmentation efficiency. The triply charged precursor of the synthetic *N*-glycopeptide E<u>N</u>YSVFVHPK carrying the NaNa *N*-glycan [M + 3H]^3+^ = 1142.12 did not yield any reasonable information on the peptide (Figure 1-A), while the quadruply charged precursor [M + 4H]^4+^ = 856.86 provided a sufficiently comprehensive z^.^ ions series (z^.^_3-8_ as singly charged ions and z^.^_9_ as doubly charged ion) that did allow peptide identification and assignment of the site of modification (Figure 1-B). The mass difference of 2318.82 Da between z^.^_8_ and z^.^_9_ indicated the presence of a glycan corresponding to the composition Hex_5_HexNAc_4_NeuAc_2_ attached to Asn, confirming the site of glycosylation within the peptide sequence. A similar charge state dependent trend was observed for the *N*-glycopeptides carrying the glycosylation site in the middle and close to the C-terminus (Supplementary Figure S1&2). Overall, we observed that the N-terminally glycosylated peptides gave balanced distribution and continuous stretches of both c- and z ions across the MSMS spectra (Supplementary Figure S1-A). These were not as prominent for the glycopeptides that carried the *N*-glycan modification in the middle or the C-terminus (Supplementary Figure S1-B and C).

**Figure 1:**
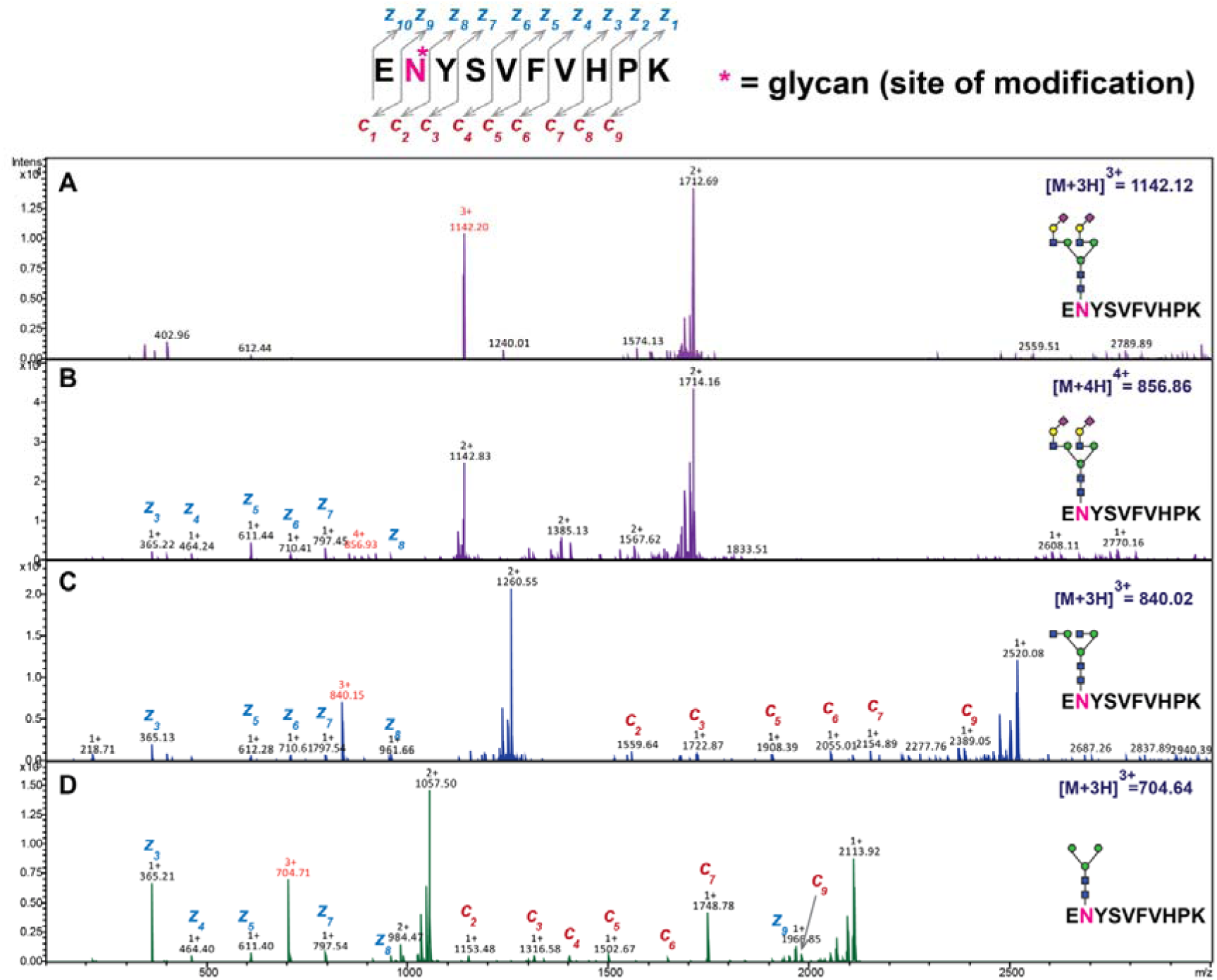
Impact of charge state and glycan size on ETD-fragmentation efficiency. ETD MS/MS spectra of a synthetic N-glycopeptide that carries the glycosylation site close to N-terminus (E**<u>N</u>**YSVFVHPK). Three different glycoforms of this glycopeptide were analysed carrying NaNa (A-B), GnGn (C) and M3 N-glycans (D, for N-glycan abbreviations please see Table 1). The precursor m/z had a significant impact on the amount of information that could be obtained on the glycopeptide backbone by ETD fragmentation. A tremendous difference in the number of detected fragment peaks was detected between the MS/MS spectra obtained for the NaNa N-glycan carrying glycopeptide when the [M + 3H]^3+^ = 1142.12 (A) or the [M + 4H]^4+^ = 856.86 (B) precursor ion were analysed. Glycopeptides with smaller sized N-glycans did yield a significantly larger number of detected c and z ions (B-D), indicating that glycan size also impacts ETD-fragmentation efficiency.

Next, the influence of the glycan size on ETD fragmentation was evaluated using the synthetic *N*-glycopeptides E<u>N</u>YSVFVHPK carrying a GnGn or M3 *N*-glycan (Table 1). In both cases, the triply charged precursors [M + 3H]^3+^ = 840.02 and [M + 3H]^3+^ = 704.64, respectively, were selected for the subsequent MSMS analyses as the quadruply charged precursors were not detected. In contrast to the NaNa carrying glycopeptide sufficient data on the c- and z^.^ ion series was obtained from the triply charged precursors to confirm the peptide backbone and site of glycosylation (Figure 1-C and D, Supplementary Figure S3-A-C). Based upon the number of detected c- and z-series ions, ETD fragmentation efficiency of glycopeptides was depending on the precursor *m/z* and glycan size, notwithstanding the fact that highly charged precursor ions (≥ 3+) were necessary for efficient fragmentation. In agreement with previous reports, we also observed that the precursor charge state had a considerable impact on ETD fragmentation [18]. Under conventional electrospray conditions most *N-*linked glycopeptides are detected in the *m/z* range above 900, which is less suited for successful ETD glycopeptide fragmentation of these compounds. This can at least partially be overcome by using charge-increasing additives such as *m*-nitrobenzyl alcohol to supercharge the analyte [19] or by the use of improved ionisation devices such as CaptiveSpray nanoBooster™ to enhance glycopeptide ionisation and increase the charge state of the analyte.

### Sialic acids negatively impact glycopeptide ETD-fragmentation efficiency

Electron capture by multiply charged peptide cations in ETD primarily results in cleavage of the N–Cα bond resulting in c′-ions and z^.^-ions. The loss of a hydrogen atom upon electron capture is well-known, resulting in a hydrogen-rich, charge reduced intact species that does, however, not fragment (no dissociation pathway, ET,noD) [5, 18, 20]. This phenomenon is more pronounced for sialic acid containing glycopeptides at low charge states. Nevertheless, the observed charge state effect on ETD fragmentation of sialylated glycopeptides likely also depends on the sites of protonation within the glycopeptide [21], relative propensities for competitive cleavage reactions [18] and possibly on the strength of electrostatic repulsion of the reactant anion by sialic acids.

The observed decreased ETD-fragmentation efficiency for sialylated glycopeptides appears to be independent on the type of glycosylation, as it has been observed for *N*-glycopeptides (Figure 1) as well as *O*-glycopeptides [22]. We observed a similar charge state dependency when further analysing human IgG 1-4 synthetic glycopeptides carrying NaNa *N*-glycan (Figure 2 and Supplementary Figure S4). Windwarder *et al* were able to overcome these undesirable, sialic acid dependant effect using enzymatic desialylation, which allowed them to sequence and identify the sites of *O*-glycosylation in c/MAM-domain of neuropilin-1 [22]. Sialic acid removal was crucial to obtain this information, with the caveat of losing site-specific glycosylation information. Nonetheless, the fact that sialic acids negatively impact ETD fragmentation needs to be considered for any glycoprotein site-specific microheterogeneity characterisation experiments based on ETD MSMS as sialic acid containing glycoforms might be underrepresented in the data due to the lack of successful identifications.

**Figure 2:**
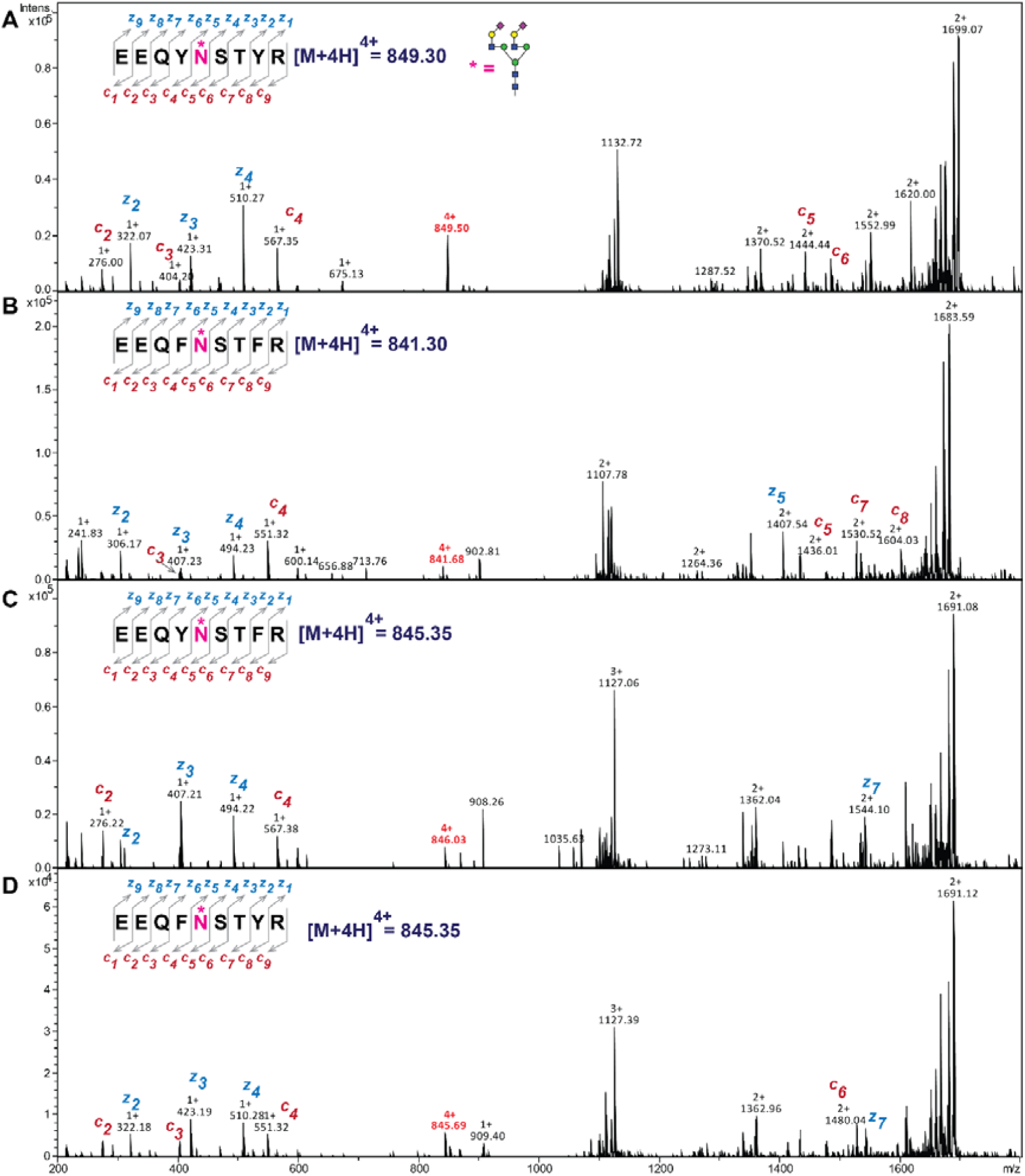
Impact of charge state and glycan size on ETD-fragmentation efficiency. ETD MS/MS spectra of synthetic IgG 1-4 (panel A-D) N-glycopeptides carrying NaNa glycan. Consistent with our previous observation, the precursor charge state had a significant impact on ETD fragmentation efficiency of sialic acid containing glycopeptides (see supplementary Figure S4 for ETD MS/MS spectra of triply charged precursor).

### Glycosylation site position within a glycopeptide impacts ETD-fragmentation efficiency

A small library of glycopeptides was used to study the effect of the glycosylation position on ETD-fragmentation efficiency on a given peptide sequence (Figure 3). The overall amino acid composition as well as the N- and C-terminal amino acids were retained, but the *N*-glycosylation sequon position was changed to obtain an N-terminally glycosylated peptide (Asn2), one carrying the *N-*glycan towards the middle (Asn7) and one immediately next to the C-terminal Lys (Asn9, Table 1). All glycopeptides carried the chitobiose core pentasaccharide (M3) as an *N*-glycan. Due to the smaller sized *N*-glycan the triply charged precursor ions were selected for ETD fragmentation as no reasonable amounts of quadruply charged signals were detected. The data suggested that the position of the glycosylation site within the sequence influenced the detected length of a continuous stretch of c- and z^.^ ions (Figure 3). The presence of a *N*-glycan pentasaccharide modification near the peptide N-terminus resulted in numerous c and z^.^ ions that were observed in the conveniently detectable *m/z* range between 400-1500 Da, whereas the presence of the same modification near the peptide’s C-terminus or towards the middle of the sequence pushed these ions further out to the border regions of the ideal MS-scan range, at least on the used instrument. This effect, however, poses a limit on what fragment ions can efficiently be trapped following fragmentation and subsequently detected to provide a sufficient stretch of diagnostic c and z ions for peptide sequence determination. In this particular example the ETD MSMS spectra provided useful data to locate the site of modification irrespective of the position of the glycosylation site. Nevertheless, this is mostly a consequence of the modest *N*-glycan size, the lack of sialic acid residues and the overall small peptide length (see also next paragraph).

**Figure 3:**
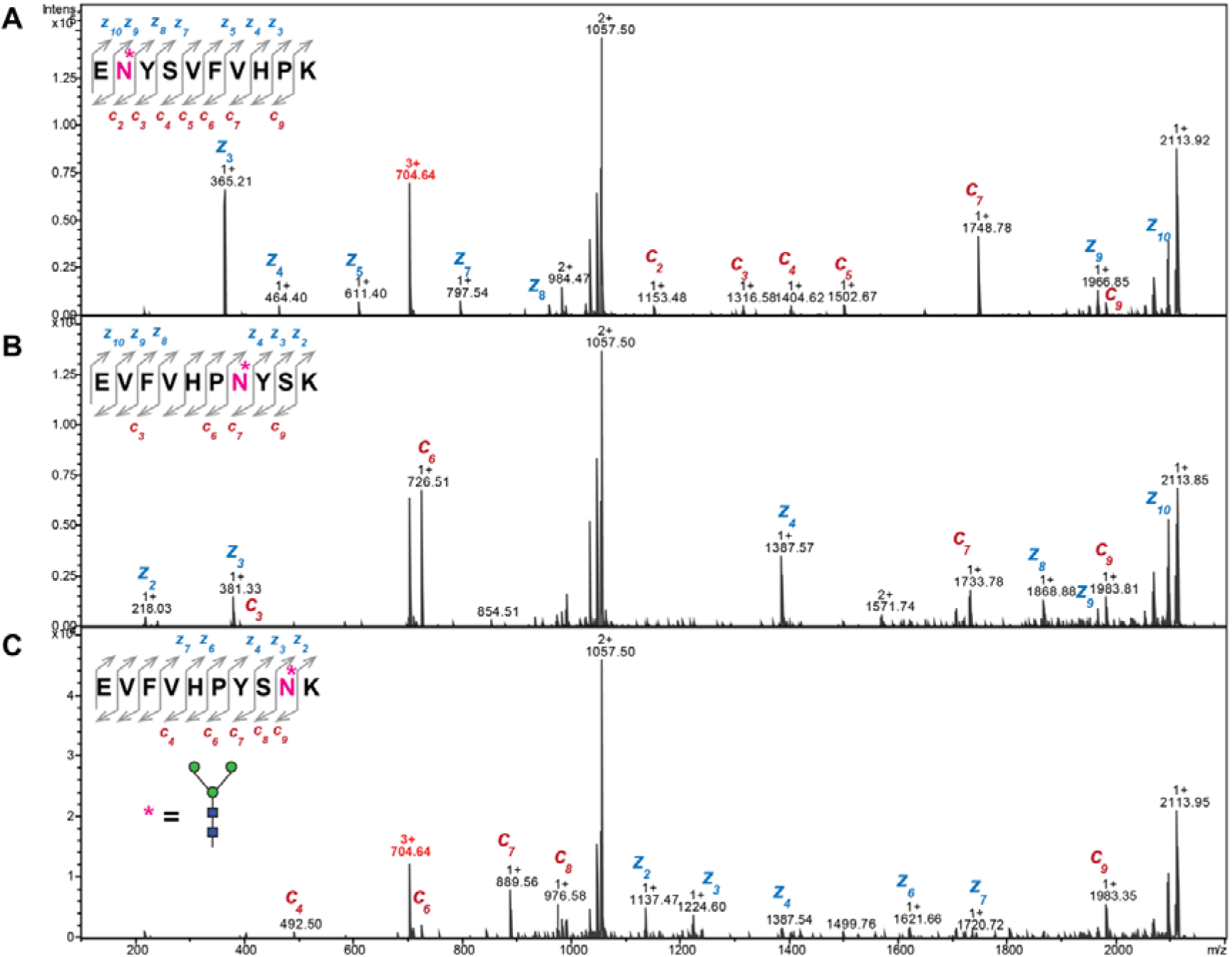
Influence of the glycosylation site location on ETD fragmentation. ETD MS/MS spectra of three isobaric glycopeptides all carrying the chitobiose core pentasaccharide N-glycan, but differing in the glycosylation site position: (A) N-terminal, (B) Middle and (C) C-terminal. The position of the glycosylation site within a peptide sequence significantly impacted ETD fragmentation efficiency and the number of detected c and z ions, which subsequently build the basis for any unambigous peptide identification.

### Glycan size and glycosylation site location matter in software assisted glycopeptide identification

Next, we tested how a conventional proteomics search algorithm such as SEQUEST and Amanda can successfully identify glycopeptides using these ETD-data sets when using defined PTM offsets. In a first step, the obtained ETD fragmentation spectra obtained for the synthetic glycopeptides were searched against a custom database containing the three peptides using Proteome Discoverer 2.1.1.21, where a set glycan modification was considered as a possible variable modification (Supplementary Table S2). The Xcorr and Amanda Score was used as an indicator for successful glycopeptide identification (Figure 4 and Supplementary Table S3). Irrespective of the glycosylation site position no successful identification was achieved from the triply charged precursor ions of the NaNa *N*-glycan carrying glycopeptides when using the Amanda search engine. Interestingly, SEQUEST was able to identify these glycopeptides. The glycopeptides carrying the M3 or the GnGn *N*-glycans, however, were successfully identified from the triply and quadruply charged precursors by both search engines (Figure 4). The highest scores were obtained for the N-terminally glycosylated glycopeptide carrying the M3 *N*-glycan, followed by the GnGn and NaNa *N*-glycopeptides. These results suggested that glycan size and location of the modification within the peptide sequence impacted how the observed peptide fragment ions were evaluated by Xcorr and Amanda and hence, the final scores given by these algorithms (Figure 4). Implementation of a scoring system that is adapted to consider different precursor charge states and peptide sequences for the analysis of ETD data can bear the potential to significantly increase the number of positively and accurately identified glycopeptides. Such an approach has already been successfully used to identify peptides using protein prospector resulting in the identification of 80% more spectra at a 1% false discovery rate [23]. It remains to be seen if and how this information can be incorporated to improve glycopeptide identification from ETD-spectra.

**Figure 4:**
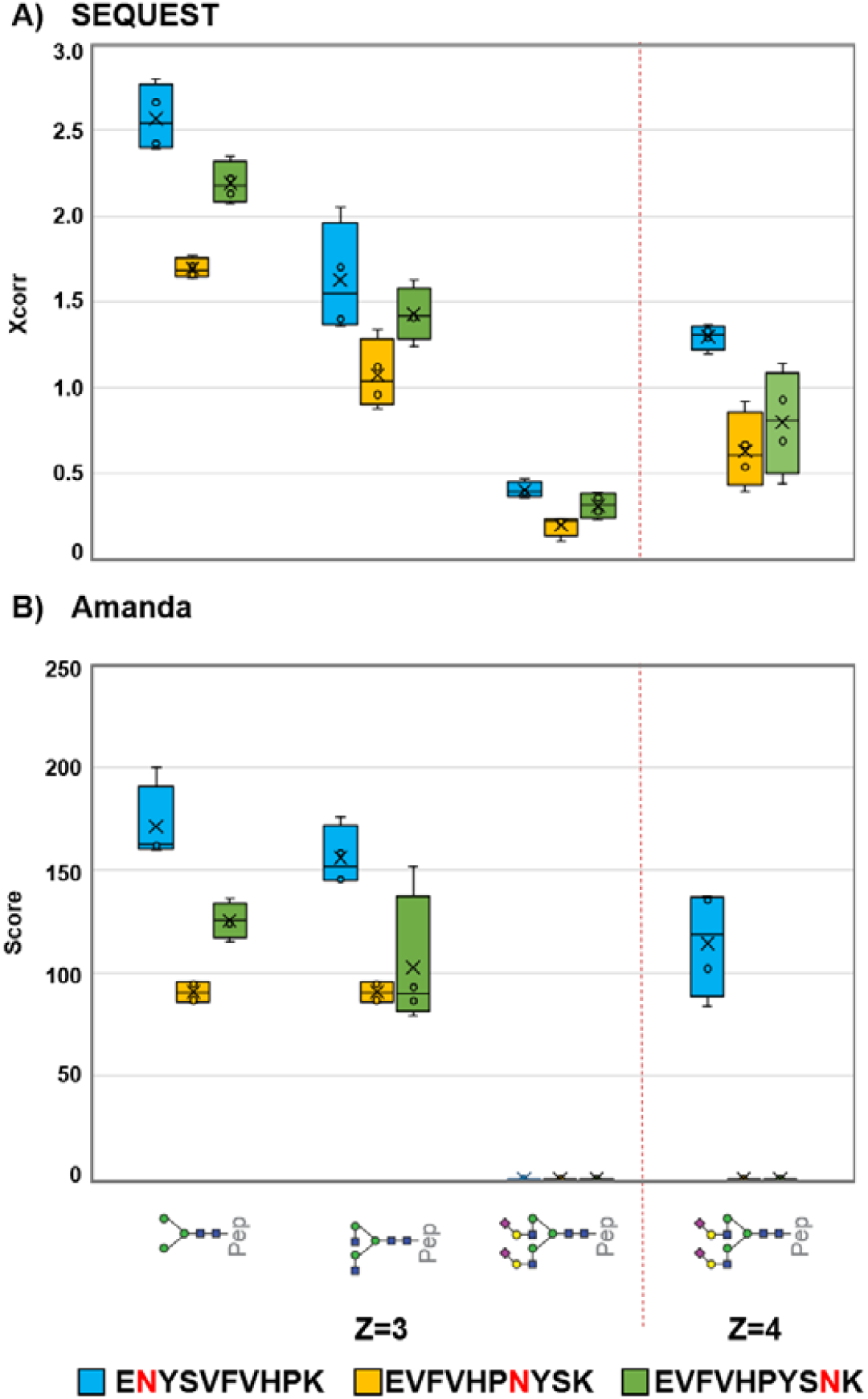
Influence of glycan size and glycosylation site location within the glycopeptide on search algorithm scores. Glycopeptides differing in N-glycan size and glycosylation site position within the peptide were analysed by LC-ESI ETD MS/MS and searched against a custom protein database using (A) SEQUEST and (B) Amanda algorithm as described in the methods section (see Supplementary Table S3).

## CONCLUSION

Our systematic study using a panel of synthetic glycopeptides provides an overview on the ETD fragmentation characteristics of glycopeptides and the associated merits & pitfalls affecting automated glycopeptide identification. Using a panel of defined, synthetic *N*-glycopeptides we evaluated how glycan size, glycosylation position within a peptide, as well as the overall charge state of the glycopeptide influenced ETD fragmentation efficiency. Our data suggest that the number and quality of assignable peptide backbone fragments was significantly depending on glycan size and the position of the modification within a peptide sequence. Peptides carrying large, sialylated glycans gave the worst score. Based upon the glycopeptide identification from ETD data using SEQUEST, and MS Amanda, we want to emphasise on the importance of the precursor charge state and *m/z* range to obtain efficient ETD fragmentation. Highly charged glycopeptides (z>3) with precursor masses of *m/z* <900 resulted in significantly more informative ETD fragment spectra. The glycan size and presence of sialic acids both negatively impacted ETD fragmentation efficiency. Improving our understanding how glycan size/composition impacts ETD fragmentation is an important step towards facilitating automated glycopeptide identification in large-scale, ETD-based glycoproteomics studies where site-assignment is crucial, such as for *O*-glycopeptides.

## ACKNOWLEDGEMENTS

We thank the *Beilstein-Institut* for supporting KA with a PhD scholarship and the Max Planck Society for financial support.

