## Supplementary Figure S1 for "Glycan size and attachment site location affect electron transfer dissociation (ETD) fragmentation and automated glycopeptide identification"

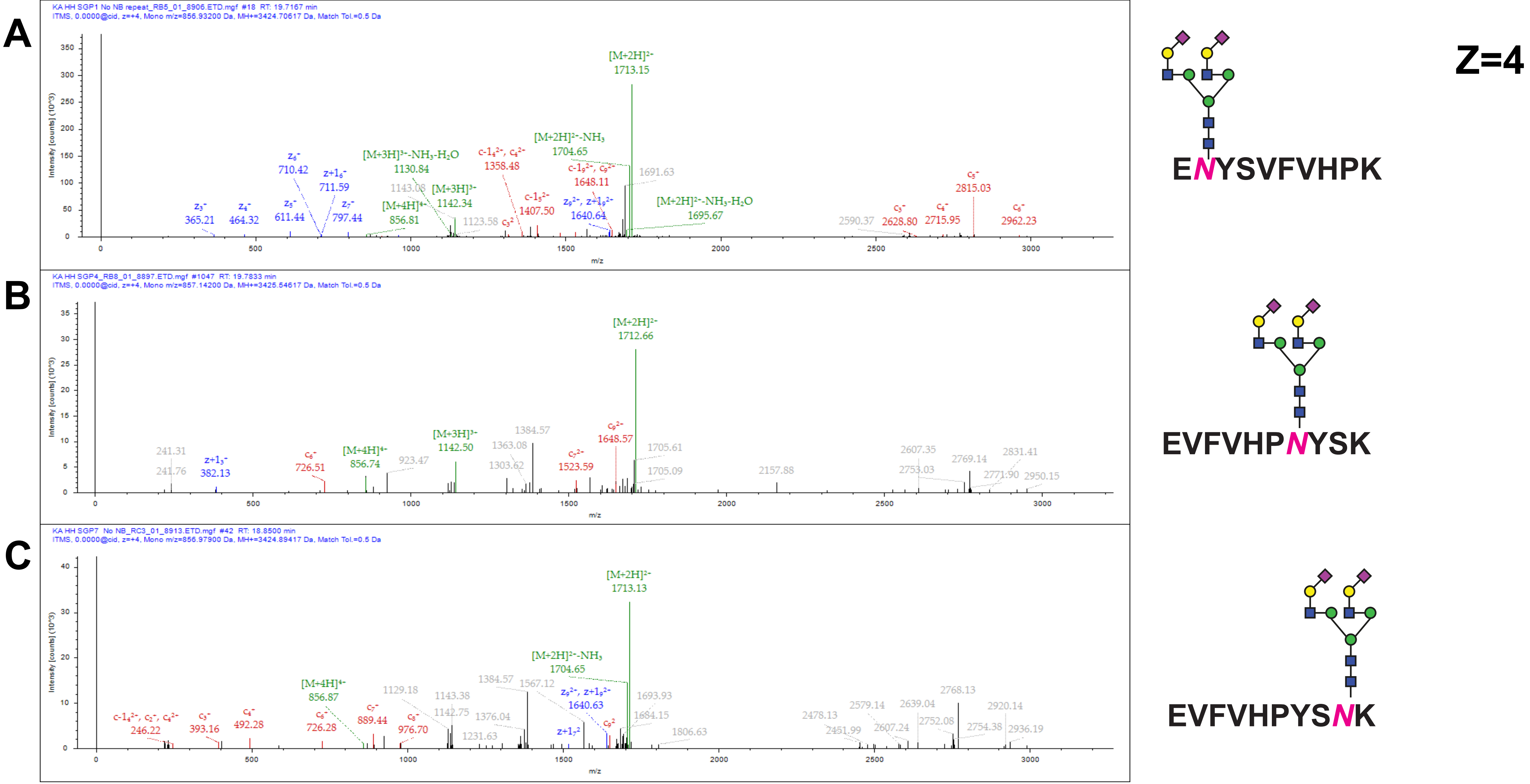


*Supplementary Figure S1: ETD MS/MS spectra for quadrply charged precursor ion of a synthetic* N-*glycopeptide that carries the glycosylation site in the (A) N-terminal, middle (B) and close to C-terminus (C) both glycopeptides carry a NaNa glycan.*

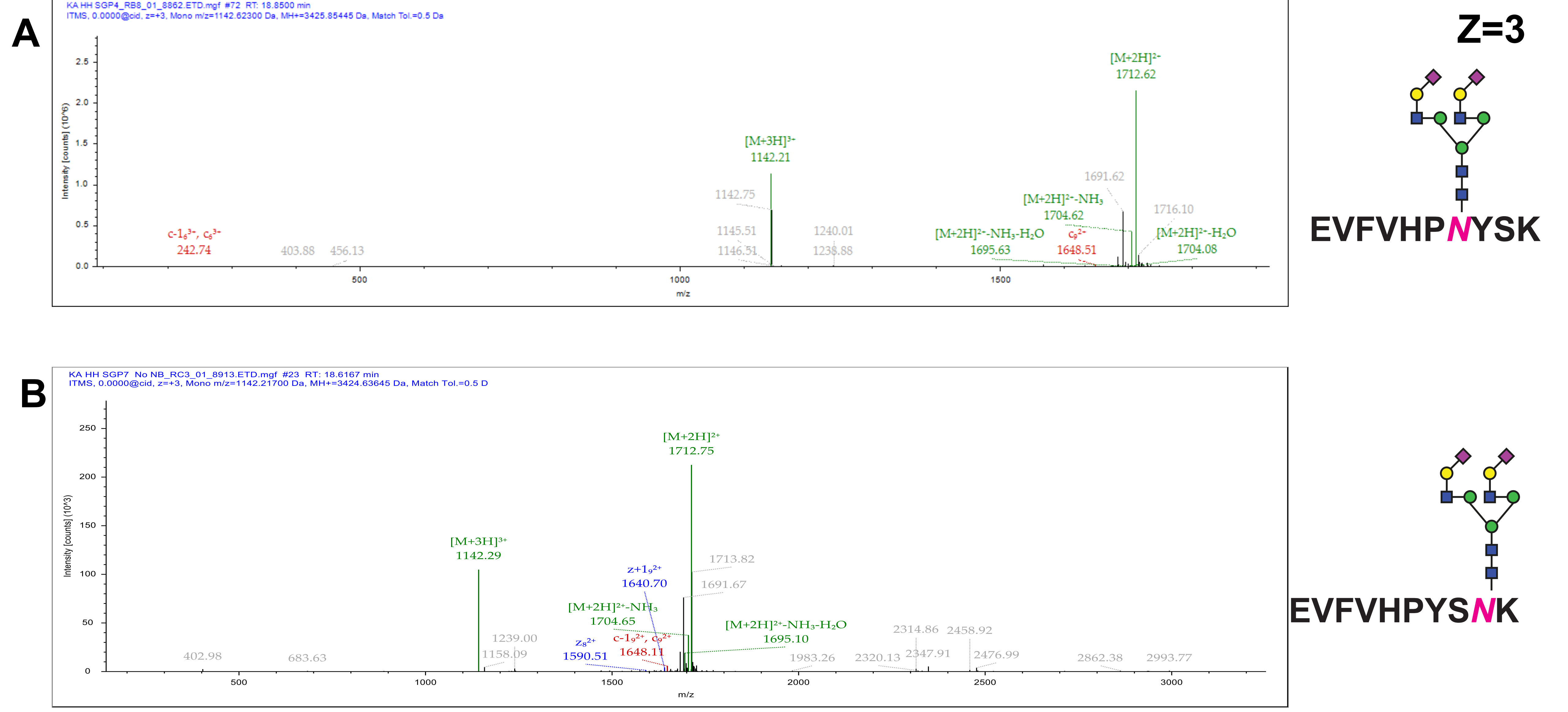


*Supplementary Figure S2: ETD MS/MS spectra for triply charged precursor ion of a synthetic* N*-glycopeptide that carries the glycosylation site in the middle (A) and close to C-terminus (B) both glycopeptides carry a NaNa glycan.*

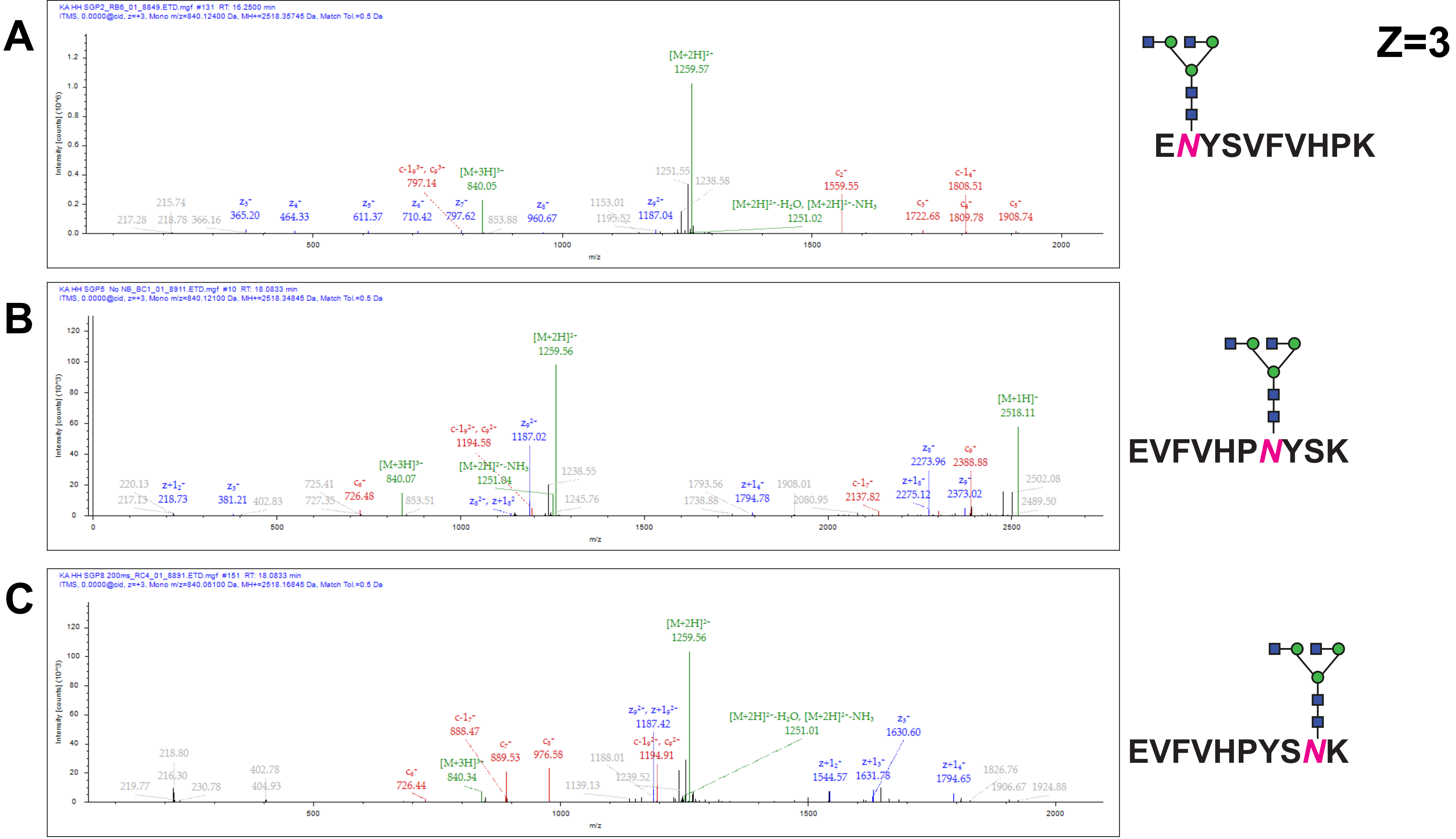


*Supplementary Figure S3: ETD MS/MS spectra obtained from the triply charged precursor ion of a synthetic* N-*glycopeptide carrying a GnGn glycan at various position (A- N-terminal, B-middle and C- C-terminal).*

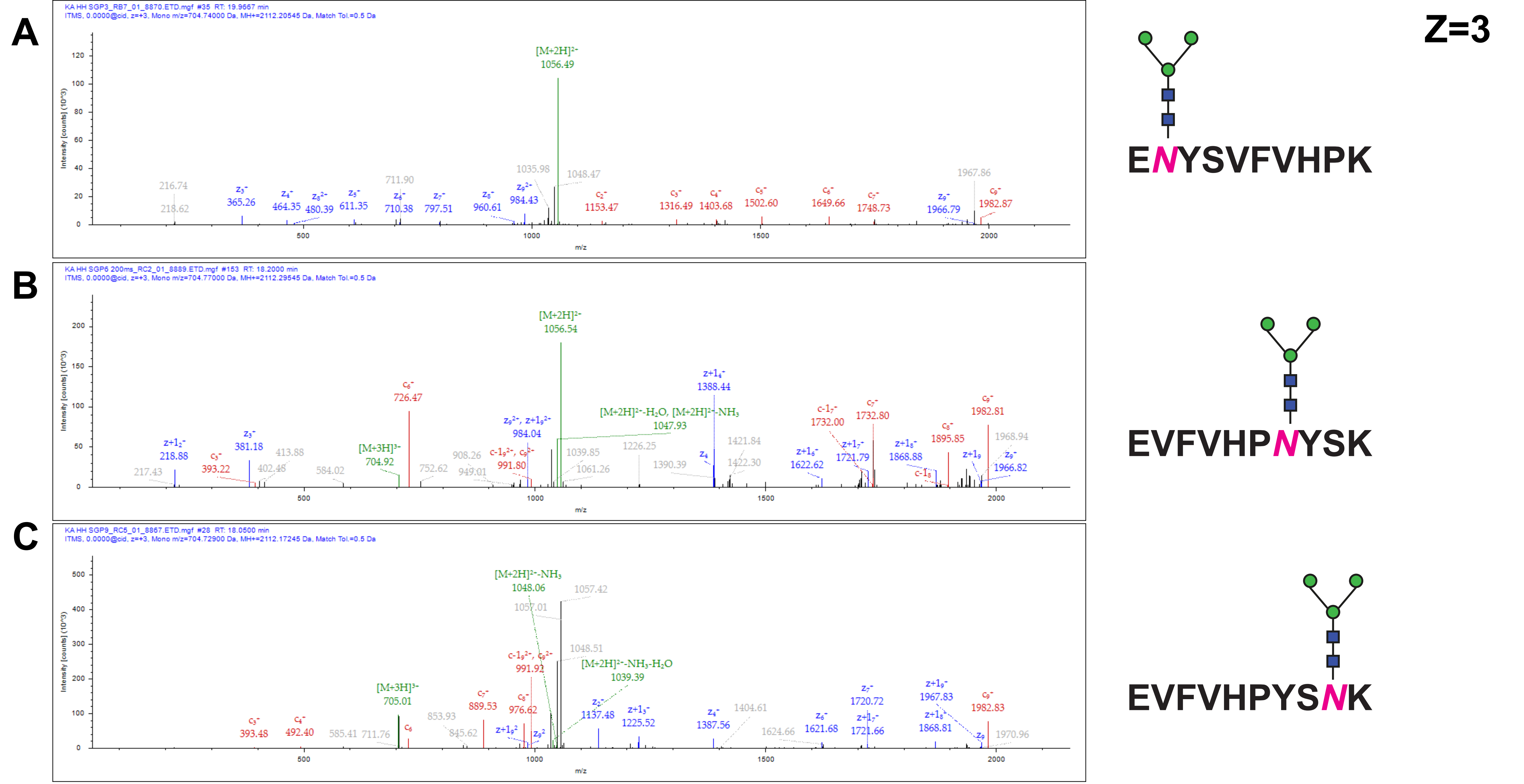


*Supplementary Figure S4: ETD MS/MS spectra obtained from the triply charged precursor ion of a synthetic* N-*glycopeptide carrying Man3 glycan at various positions (A: N-terminal, B: middle and C: C-terminal).*

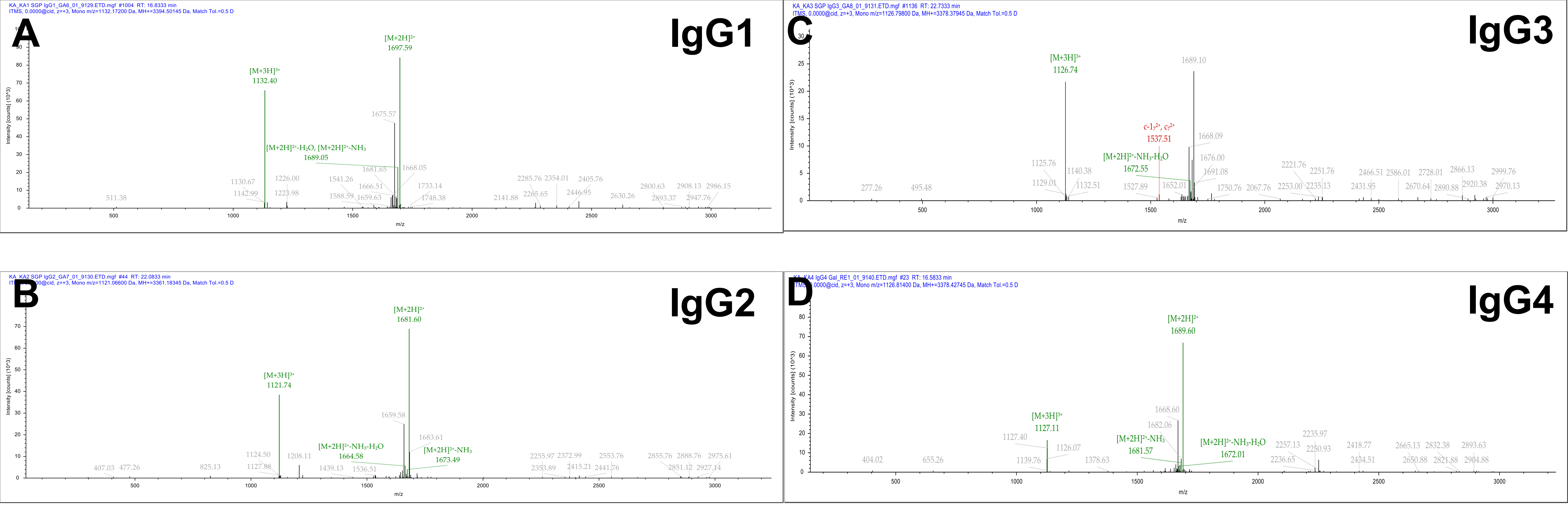


*Supplementary Figure S4: ETD MS/MS spectra obtained from the triply charged precursor ions of synthetic IgG 1-4* N-*glycopeptides carrying the* NaNa N-*glycan from triply charged precursor ion (A) IgG 1 (B) IgG 2 (C) IgG 3 (D) IgG 4*
